# Biological insights from the whole genome analysis of human embryonic stem cells

**DOI:** 10.1101/2020.10.26.337352

**Authors:** Florian T. Merkle, Sulagna Ghosh, Giulio Genovese, Robert E. Handsaker, Seva Kashin, Konrad Karczewski, Colm O’Dushlaine, Carlos Pato, Michele Pato, Daniel G. MacArthur, Steven A. McCarroll, Kevin Eggan

## Abstract

There has not yet been a systematic analysis of hESC whole genomes at a single nucleotide resolution. We therefore performed whole genome sequencing (WGS) of 143 hESC lines and annotated their single nucleotide and structural genetic variants. We found that while a substantial fraction of hESC lines contained large deleterious structural variants, finer scale structural and single nucleotide variants (SNVs) that are ascertainable only through WGS analyses were present in hESCs genomes and human blood-derived genomes at similar frequencies. However, WGS did identify SNVs associated with cancer or other diseases that will likely alter cellular phenotypes and may compromise the safety of hESC-derived cellular products transplanted into humans. As a resource to enable reproducible hESC research and safer translation, we provide a user-friendly WGS data portal and a data-driven scheme for cell line maintenance and selection.

**GRAPHICAL ABSTRACT:** 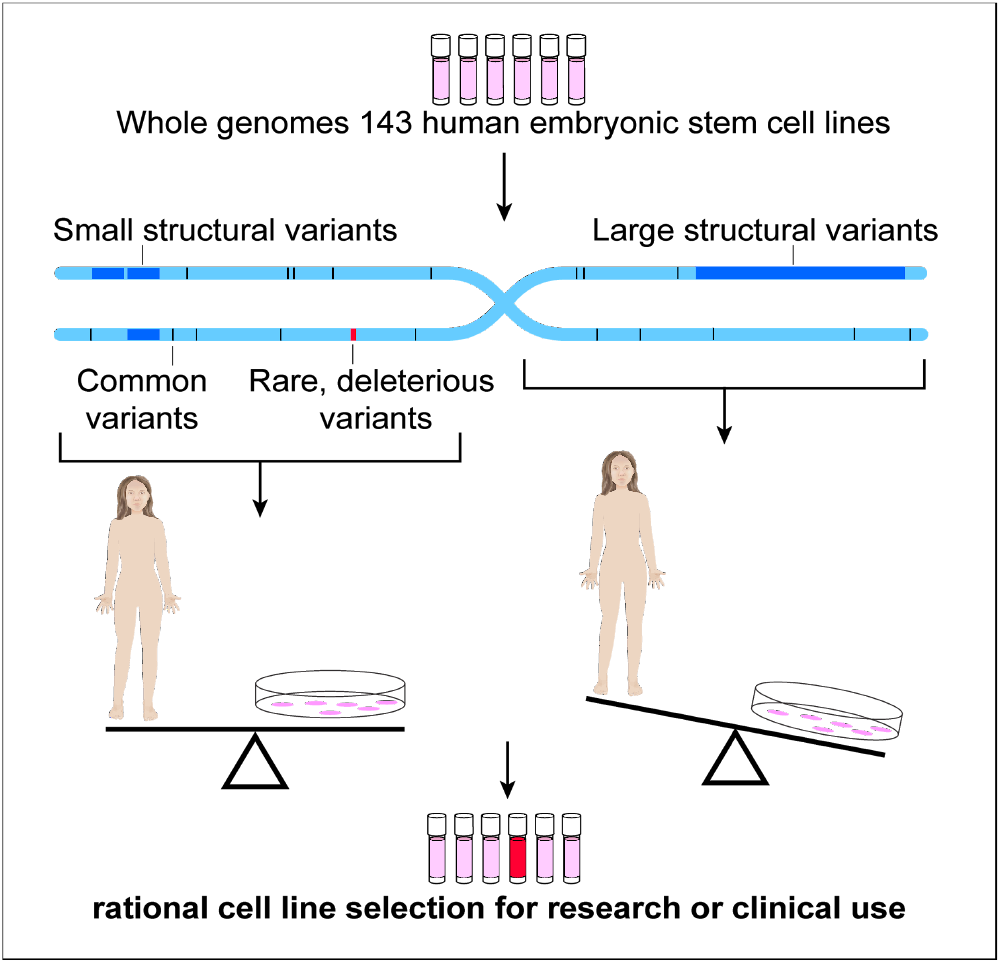

**IN BRIEF:** Merkle and Ghosh et al. describe insights from the whole genome sequences of commonly used human embryonic stem cell (hESC) lines. Analyses of these sequences show that while hESC genomes had more large structural variants than humans do from genetic inheritance, hESCs did not have an observable excess of finer-scale variants. However, many hESC lines contained rare loss-of-function variants and combinations of common variants that may profoundly shape their biological phenotypes. Thus, genome sequencing data can be valuable to those selecting cell lines for a given biological or clinical application, and the sequences and analysis reported here should facilitate such choices.

**HIGHLIGHTS:**

- One third of hESCs we analysed are siblings, and almost all are of European ancestry
- Large structural variants are common in hESCs, but finer-scale variation is similar to that human populations
- Many strong-effect loss-of-function mutations and cancer-associated mutations are present in specific hESC lines
- We provide user-friendly resources for rational hESC line selection based on genome sequence

## INTRODUCTION

Human pluripotent stem cells (hPSCs) can self-renew indefinitely while retaining the ability to differentiate into many cell types (Jones and Thomson, 2000). These properties make hPSCs a powerful resource for studying early human development, disease modelling, drug discovery, and increasingly for developing candidate cell therapies (Avior et al., 2016; Merkle and Eggan, 2013; Sandoe and Eggan, 2013; Trounson and DeWitt, 2016) (https://clinicaltrials.gov). However, the utility of hESCs and human induced pluripotent stem cells (hiPSCs) for these applications can be compromised by mutations that affect their differentiation potential, cellular phenotypes, or clinical safety. The nature of such mutations has been studied using karyotyping (Draper et al., 2004; Lund et al., 2012), fluorescent in situ hybridization and comparative genome hybridization arrays (Dekel-Naftali et al., 2012; Spits et al., 2008), and high-density single nucleotide polymorphism (SNP) DNA microarrays (Amps et al., 2011; Laurent et al., 2011; Lefort et al., 2008; Närvä et al., 2010). Despite the limited spatial genomic resolution of these methods (typically > 100 kbp), these studies revealed recurrent, culture-acquired structural variants including duplications of chromosomes 1, 12, 17, 20, X, or segments thereof. While the selective advantage conferred by one of these recurrent duplications at Chr20q11.21 has been attributed to the gain of the anti-apoptotic gene *BCL2L1* (Avery et al., 2013; Nguyen et al., 2014), the cause and functional consequences of most mutations observed in hPSCs remain poorly understood.

Despite their considerable use in basic research and their increasing use in the development of cell therapies for transplantation into humans, as much as 99% of the genome of most hPSCs remains unexplored. Because the genome sequences of most stem cell lines are poorly ascertained, research groups have had little choice but to view hPSC lines as largely interchangeable and to select lines for a given application by convenience or by historical precedence rather than their intrinsic genetic suitability (Kobold et al., 2015; Löser et al., 2010). The lack of comprehensive genetic information about cell lines of interest is also troubling since mutations can accumulate in hPSCs over time in culture (Laurent et al., 2011), potentially compromising experimental reproducibility or safety. Establishing a baseline whole genome sequence for as many hESC lines as possible would allow such culture-acquired mutations to be more easily identified in the future (Avior et al., 2019).

To start addressing these important issues, we (Merkle et al., 2017) and others (Kilpinen et al., 2017) recently analysed the whole exome sequences (WES) of hPSCs. We found many genetic variants that had likely been acquired in culture and discovered that approximately 5% of widely used cell lines had acquired mutations in the tumour suppressor gene *TP53* (p53) (Merkle et al., 2017). These mutations, which were confirmed by others (Amir et al., 2017; Avior et al., 2019), conferred significant growth advantage in culture but did not prevent differentiation to multiple cell lineages.

Given that exome sequencing many hESC lines could reveal important biological insights, we wondered what might be learned by examining their whole genomes. We therefore performed whole genome sequencing (WGS, >25x coverage) and complementary high-density SNP genotyping of 143 hESC lines and report our findings here as a resource. We found that the overall burden of both SNVs and CNVs in hESCs resembled that of human populations, indicating that almost all variants were inherited and thus confirming hESCs as invaluable tools for studying human biology. However, our analyses also bring to light previously unreported recurrent acquired genetic variants that point to selective pressures exerted during self-renewal. These included a recurrent amplicon on Chr1q32.1, a copy-neutral loss of heterozygosity (CN-LOH) event at Chr9q, and small deletions encompassing the gene *EP300*, whose gene product stabilizes p53. Additional studies of SNVs we identified from WGS data revealed likely inherited and deleterious variants in genes associated with cancer, infertility, and a variety of autosomal dominant diseases that will likely impact the phenotypic behaviour of individual stem cell lines that harbour them. In order to allow any researcher to query our data and to facilitate rational cell line selection for a given application, we developed a user-friendly online data portal, which will further the goals of experimental reproducibility and the safety of future cell therapies.

## RESULTS

### HESC line selection and whole genome sequencing

To gain insight into stem cell biology and generate a valuable resource for the research and medical communities, we sequenced the whole genomes of 143 hESC lines that had been voluntarily deposited into the registry of hESCs approved for federally funded research that is maintained by the US National Institutes of Health (NIH) (http://grants.nih.gov/stem_cells/registry/current.htm) or that had been prepared for therapeutic applications (Fig. 1). Genomic DNA from these cell lines was sequenced to a mean read depth of 32.2 (SD 6.4, range 23.3 for HUES68 to 60.9 for KCL038), (Fig. S1A) resulting in an average of 97% of the genome being sequenced at a minimum of 10x coverage (Fig. S1B). Sample information is detailed in Table S1A, and sequencing and analysis methods are described in Materials and Methods.

**Figure 1.**
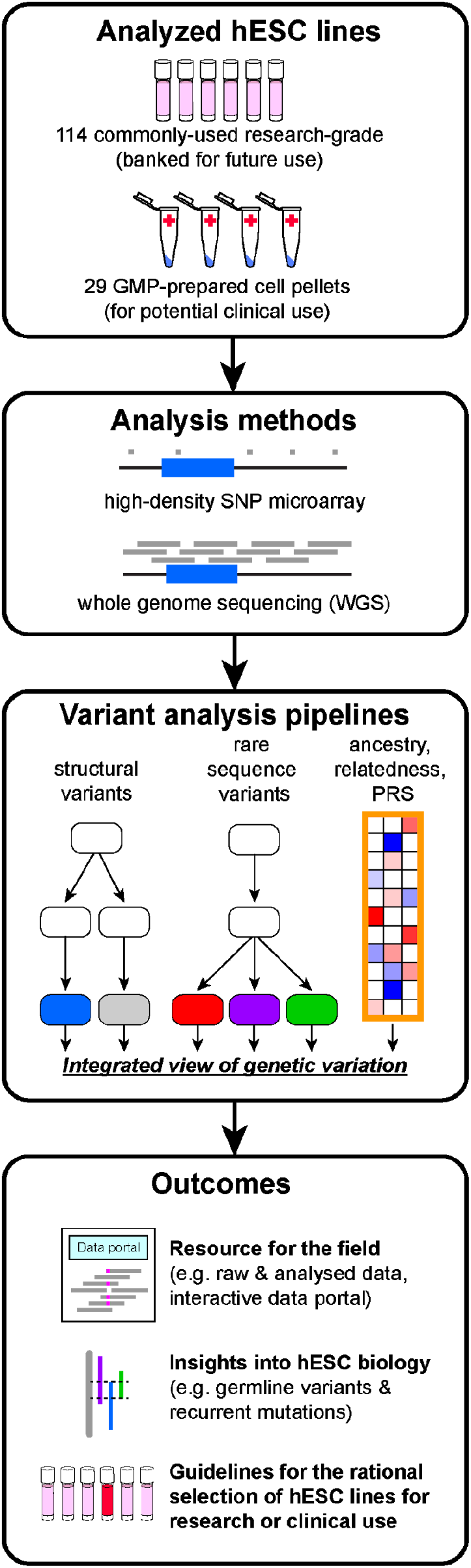
Study design and outputs. The genomic DNA of 143 human embryonic stem cell lines (hESCs) was analyzed by high-density SNP microarray and at the single-nucleotide level by whole genome sequencing to call structural variants to a resolution of ~1 kbp, rare sequence variants associated with disease, and common sequence variants to reveal cell line ancestry, relatedness, polygenic risk score (PRS). An integrated analysis of these data, which are provided as a resource to the field via an interactive data portal, yields novel insights into hESC biology and facilitates the rational selection of cell lines based on their genetic architecture.

### hESCs are predominantly European and often share sibling relationships

Genetic background can be an important modifier of cellular phenotypes (Rouhani et al., 2014a; Sittig et al., 2016). We therefore first investigated the genetic ancestry of hESC lines by drawing upon their ancestry-informative SNPs and comparing them to the diverse human populations sequenced in the 1000 Genomes Project (Consortium et al., 2012, 2015). In agreement with previous studies (Mosher et al., 2010), principal component analysis (PCA) revealed that 93% (133/143) of sequenced hESC lines clustered together with samples of European ancestry (Fig. 2A,B) and predominantly with samples of Central European (CEU) or British (GBR) ancestry (Table S1B,C). A smaller number of lines (9/143) had likely admixtures of European and other ancestries, while one was of apparent East Asian ancestry (CSES12). The predominantly European ancestry of stem cell genotypes was also reflected in their human leukocyte antigen (HLA) haplotypes (Nunes et al., 2014) (Fig. S1C, Table S2A). These HLA genotypes may be useful for groups seeking to match stem cell-derived transplant and recipient HLA haplotypes (Aron Badin et al., 2019; Morizane et al., 2017).

**Figure 2.**
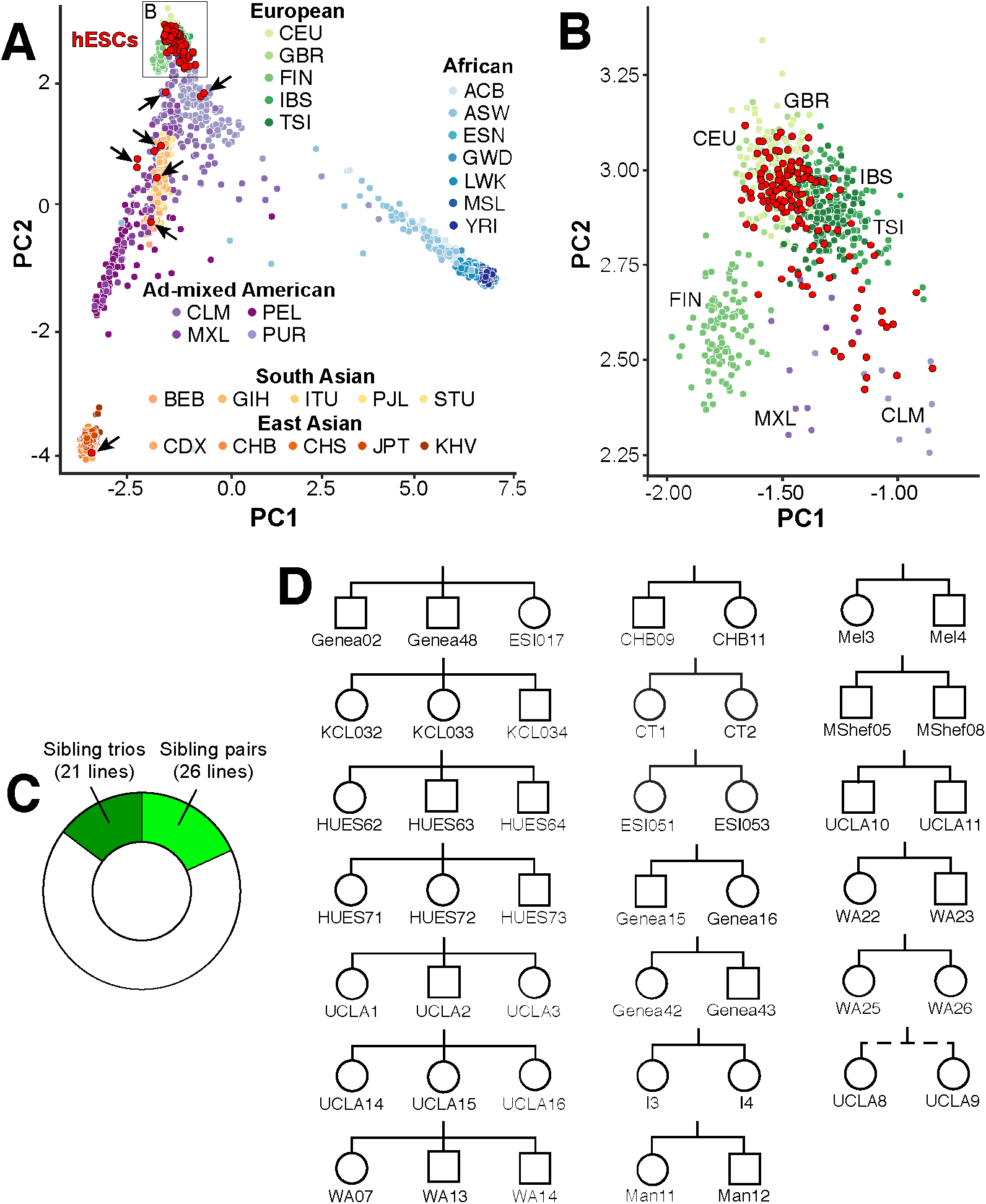
Ancestry and relatedness. **A)** Principal component analysis illustrates the genetic ancestry of hESC lines (red with black outline) relative to individuals from diverse populations (three-letter codes) from the 1000 Genomes project. HESCs with ad-mixed European or non-European ancestry are indicated by arrows. PC, principal component. **B)** Magnification of the hESC cluster from (A) shows clustering of hESC lines cluster with samples of Central European, British, and Southern European ancestry. **C)** One third of hESC lines have direct sibling relationships. **D)** Sibship pedigree of hESC lines, where squares denote male, circles denote female, and the dashed line denotes a half-sibling relationship.

Since multiple hESC lines can be derived from a cohort of embryos donated by a single couple undergoing assisted reproduction by *in vitro* fertilization (IVF) (Chen et al., 2009) we wondered how many sequenced hESC lines might exhibit genetic relatedness to one another. Upon analysing shared SNP alleles (see Materials and Methods), we found that 47/143 (33%) hESC lines shared a direct sibling relationship with another line we had sequenced, including seven sibling trios, 12 sibling pairs, and one half-sibling pair (Fig. 2C,D Table S1A-C). Importantly, many of these sibling relationships were either unknown or unreported. For example, one sibling trio contained hESC lines from distinct providers (Genea02, Genea48, ESI017) (Fig. 2D). Upon contacting the providers we learned that these cell lines were derived using materials from the same IVF clinic and that the sibling trio also included a fourth line (ESI014) not available for distribution. Awareness of these familial relationships should help guide experimental design, which in some contexts may aim to avoid shared genetic background, and in other contexts might exploit these properties to test genotype-phenotype relationships.

### Common genetic variant contribution to risk of disease phenotypes

Common single nucleotide polymorphisms (SNPs) can impact the suitability of cell lines for modelling disease or transplantation. For example, a genetic variant in the gene *ABO* causes the O blood type (Yamamoto et al., 1990), a variant in *CCR5* renders cells resistant to HIV infection (Dean et al., 1996; Samson et al., 1996), while variants in the genes *APOE* (Corder EH et al., 1993) and *TREM2* (Guerreiro et al., 2013; Jonsson et al., 2013) are among the strongest known genetic contributors to cardiovascular disease (*APOE*) and Alzheimer’s Disease (AD, both *APOE* and *TREM2*). We therefore genotyped hESCs for these common variants and identified 22 cell lines with a “universal donor” O blood type, a cell line (Elf1) likely resistant to HIV infection (Ware et al., 2014), and three cell lines homozygous for the *APOE* “e4/e4” risk haplotype for cardiovascular disease and AD (Fig. 3A-C, Table S2B).

**Figure 3.**
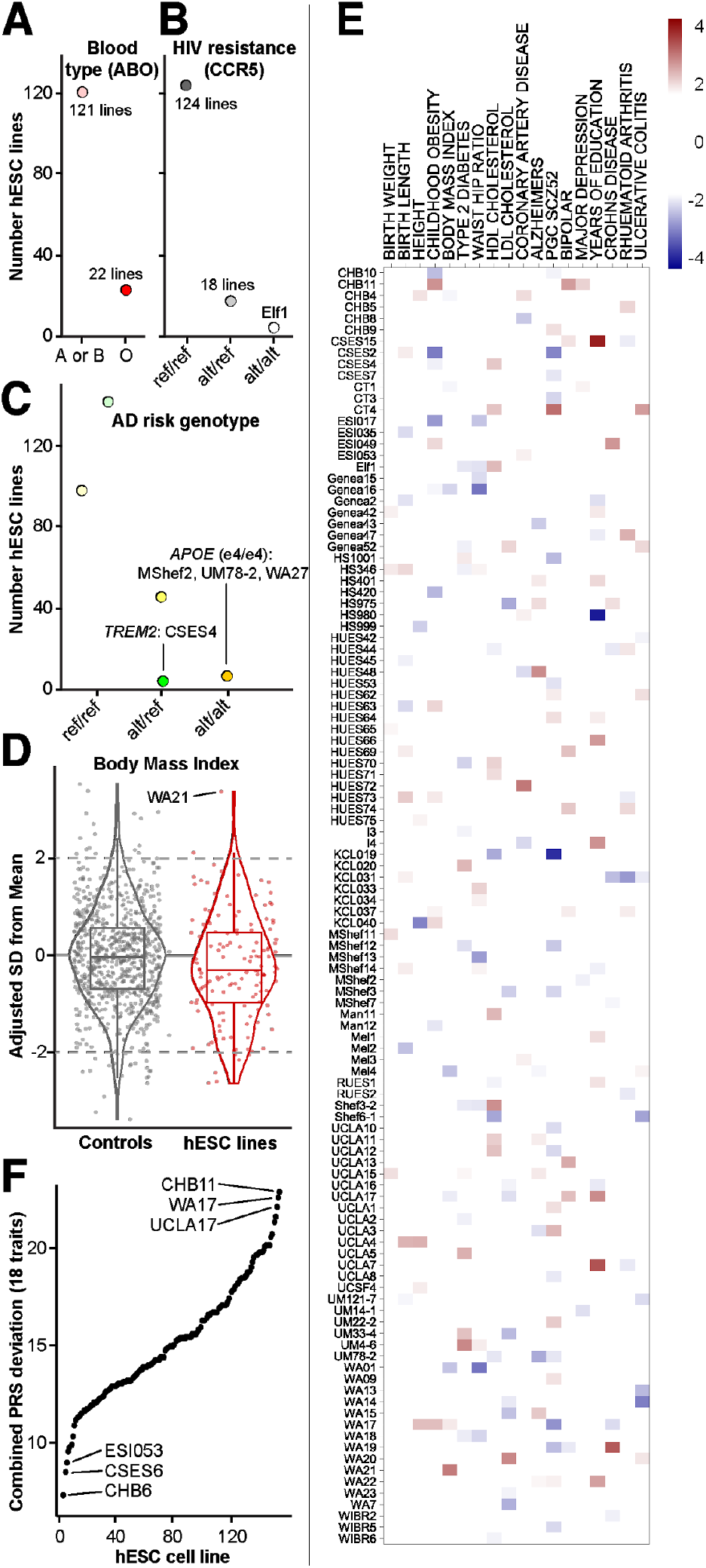
Disease risk from common genetic variants. **A-C)** Genotyping results at SNPs indicative of A) blood type (rs8176719 in *ABO*), B) resistance to HIV infection (rs333 at CCR5) and C) risk for Alzheimer’s Disease (AD, rs429358 and rs7412 in *APOE* and rs75932628 in *TREM2*). ref, reference allele; alt, alternate (risk) allele. **D)** Distribution of polygenic risk scores (PRS) for control samples and hESCs after ancestry adjustment and normalization reveals “outlier” samples with standard deviation (SD) of 2 or more. **E)** Heatmap of PRS for each of the 18 analysed traits for cell lines that were an outlier for at least one trait. **F)** Combined PRS deviation from mean for each of the 18 analysed traits.

There is also accumulating evidence that the combined actions of thousands of common SNPs can contribute substantially to risk for developing certain conditions, often conferring as much risk as large-effect (Mendelian) variants (Khera et al., 2018). The quantifiable contribution these SNPs confer can be represented through a polygenic risk score (PRS) (Khera et al., 2019). To determine the currently calculable risk conferred by such variants to distinct disease phenotypes in each cell line, we computed PRSs for 18 distinct traits using data from well-powered genome-wide association studies (GWAS) adjusted for ancestry, and normalized these scores to the distributions formed by larger numbers of similarly-sequenced human samples (Bulik-Sullivan et al., 2015b; Ripke et al., 2014) (Fig. 3D, Table S2C,D, Fig. S2). For each trait, we found one or more “outlier” hESC lines with a PRS at least two standard deviations (SD) from the mean. For example, WA21 has a high PRS for body mass index (BMI), suggesting it might be predisposed to display obesity-relevant phenotypes if differentiated to relevant cell populations such as hypothalamic neurons (Merkle et al., 2015; Wang et al., 2015). Overall, 112/143 (78%) cell lines were outliers for at least one trait, and each cell line had a unique PRS fingerprint (Fig. 3E). To identify hESCs that might make good “all-purpose control” cell lines, we ranked hESC lines by their combined absolute PRS across the 18 traits and identified cell lines with PRSs close to the population mean (Fig. 3F, Table S2E). Overall, our sequence resource will provide an opportunity to wield hESC based models to study how common genetic variants affect cellular phenotypes in a more informed manner.

### Calling structural genetic variation from WGS data

Having established the genetic background of hESCs using common SNPs, we next analysed their structural variants, which can affect the expression of tens to thousands of genes and significantly alter cellular phenotypes (Chiang et al., 2017). In particular, aneuploidy and large CNVs often contribute to disease (Henrichsen et al., 2009) and copy-neutral loss of heterozygosity (CN-LOH) events are frequently associated with cancer and can potently alter gene expression by affecting imprinted genes and unmasking disease-associated recessive mutations or risk alleles (Nicholls et al., 1989).

We reasoned that the WGS data with at least 25x mean sequencing depth of coverage we obtained should provide both superior spatial resolution and sensitivity for detecting small or mosaic CNVs in hESCs than do the data from SNP microarrays that sample only a small fraction of nucleotides (Amps et al., 2011; Canham et al., 2015). Indeed, we found that normalized read depth analysis of WGS from 121 cell lines permitted the identification of deletions as small as ~1.1 kbp and duplications as small as ~2.8 kbp (Fig. S3A). To complement this analysis, we identified heterozygous SNPs across the genome and compared the sequencing depth of both alleles in all 143 hESC lines (Alkan et al., 2011) to call CNVs and CN-LOH events. We next arbitrarily split structural variants into “large” (>1 Mbp) and “small” (<1 Mbp) categories, (Fig. S3C, Table S3A) revealing 67 distinct large structural variants affecting nearly a third of hESC lines (46/143, 32%, Table S3B).

To test the accuracy and sensitivity of our approach, we compared WGS structural variant calls to a published SNP microarray-based study (Canham et al., 2015) that included 22 of the cell lines we subsequently sequenced. We found that WGS confirmed most of these variants, allowed CNV borders to be more accurately mapped, and revealed previously unascertained structural variants (Fig. 4A, Table S3C,D). To broaden this comparison, we analysed identical genomic DNA samples from 121 hESC lines by both WGS and high-density SNP microarrays (Infinium PsychArray, > 500,000 probes), and compared the resulting structural variant calls (Table S3E). Only 27 of the 67 large variants (>1 Mbp) observed by WGS were also called by PsychChip (Fig. 4B). Together, these results suggest that analysis of WGS data has substantially improved utility for calling large structural variants relative to microarrays and confirms that hESCs carry an excess burden of large structural variants compared to somatic human cells.

**Figure 4.**
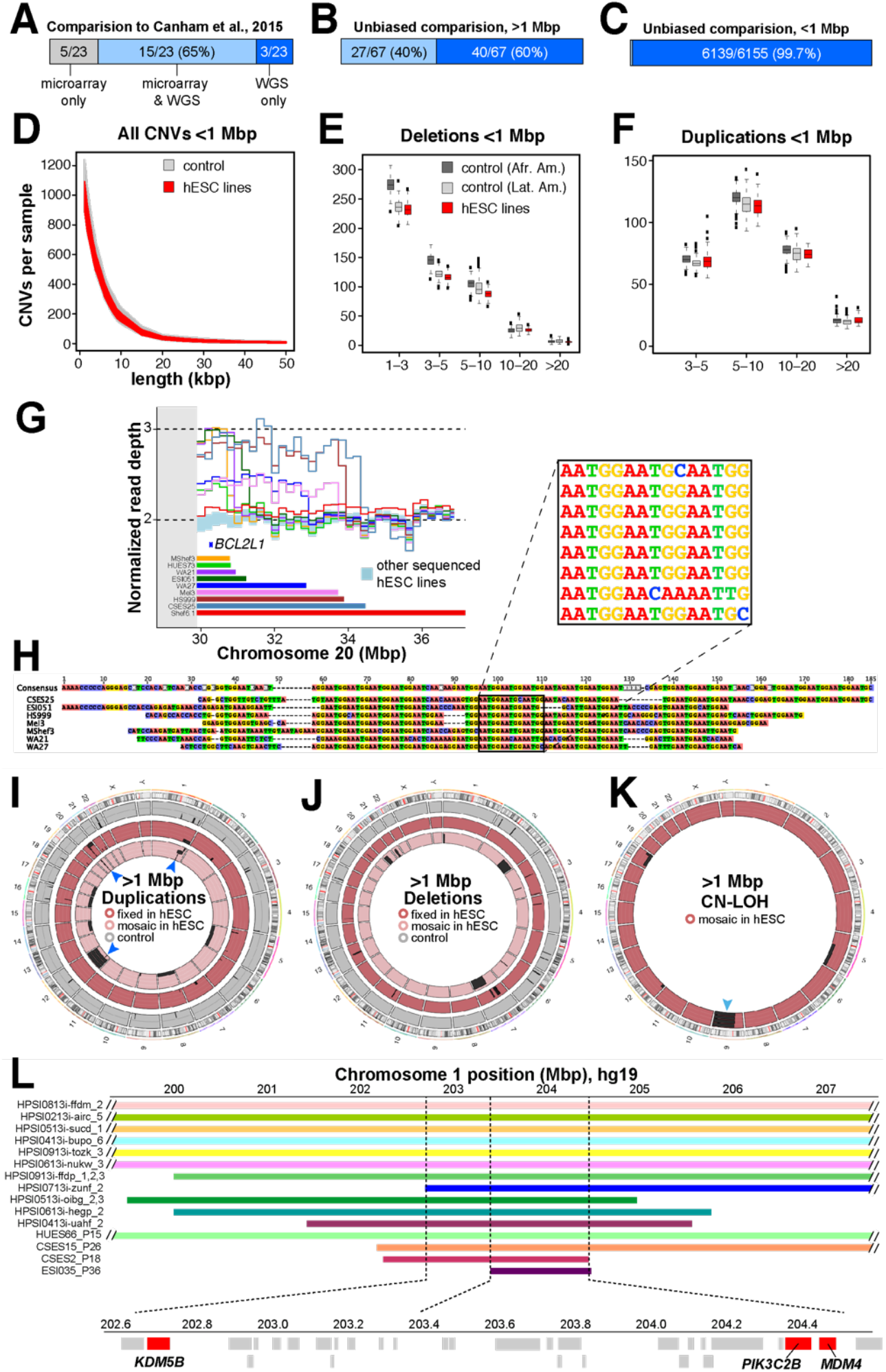
Structural variant calling from WGS data. **A)** Comparison of structural variant calls in 22 lines shared between this study and a previous publication. **B,C)** Comparison of large (B) and small (C) structural variant calls made by microarray or by WGS. **D)** Length distribution of small duplications and deletions in hESCs and controls. **E,F)** At length scales of ~1.1 kbp to 1 Mbp, the frequency of deletions (E) and duplications (F) in hESCs is indistinguishable from human primary cells. X-axis values of 1-3 and 3-5 refer to 1.117-2.75 kbp and 2.75-5 kbp, respectively. Afr. Amer., African American; Lat. Amer., Latin American. **G)** Observed recurrent duplications at the pericentromeric (grey) region of Chr20q11.21 encompassing the anti-apoptotic gene *BCL2L1*. **H)** Alignment of sequencing reads flanking the 3’ end of the Chr20q11.21 duplication reveals a shared (AATGG)_n_ motif. **I-K)** Circular ideograms of fixed and mosaic duplications (J), deletions (K) and CN-LOH events (L) in 121 hESC lines and 500 similarly-sequenced controls. Regions with at least 4 recurrent events in hESCs are indicated by blue arrowheads. **L)** A minimally duplicated region on Chr1q32.1 harbors candidate genes (red).

### Frequency of small structural variants in hESCs

Since large CNVs were relatively common in hESCs, we wondered if these stem cells might also carry an excess burden of small CNVs, which have not yet been comprehensively examined in any detail. To address this question, we filtered out genomic regions containing CNVs >1 Mbp from affected samples, and studied the residual whole genomes of 121 hESCs alongside 243 control whole genome sequences from human blood and lymphoblastoid cell line (LCL) samples (Pato et al, unpublished) for CNVs between 1.1 kbp and 1 Mbp. We observed 6155 unique CNVs, many of which were shared across cell lines, leading to an average of 999 ± 66 small CNVs per hESC sample (Table S3F,G). These numbers were in vast excess of what was observable from SNP microarrays (16/6155, 0.3%, Fig. 4C) using depth of coverage methods. However, the frequencies of these smaller deletions and duplications in hESCs, as assessed from WGS data, were indistinguishable at all length scales tested from those in similarly sequenced populations of human somatic cells (Fig. 4D-F).

Since the overall CNV burden was large, we hypothesized that it might be possible to detect a signature of culture-acquired variants in hESCs based on their acquisition of genomic regions not commonly associated with CNVs in human populations. We therefore identified the CNVs present just once in the combined dataset of hESCs and control samples. We found that each hESC line harboured on average of 21 ± 11 of these singleton CNVs, which was again indistinguishable from the frequency of similar events in the somatic cell control samples (Fig. S3D-F, Table S3G). Together, our findings are consistent with the notion that hESCs and somatic cells from many individuals carry a similar overall burden of small duplications and deletions.

### Location and potential roles of structural variants in hESCs

To gain insight into the mechanisms that contribute to recurrent structural variants in hESCs, we examined the well-studied Chr20q11.21 region (Amps et al., 2011; Spits et al., 2008) and observed duplications in 11/143 (8%) cell lines we subjected to WGS (Fig. 4G), including one additional likely instance of isochromosome 20 (Fig. S3G). The Chr20q11.21 duplication extends from regions near the centromere that cannot be accurately mapped to variable breakpoints along the long arm of Chr20. Chr20q11.21 contains the anti-apoptotic gene *BLC2L1* which has been shown to confer a selective advantage when duplicated (Avery et al., 2013; Nguyen et al., 2014). However, it remains unclear why this genomic region is so much more frequently duplicated than other regions that harbour similar anti-apoptotic genes or proto-oncogenes. To ask if the local sequence context might shed light on this issue, we mapped the non-centromeric Chr20q11.21 CNV breakpoints. These breakpoints were unique for each cell line (Fig. 4H), but all shared a common centromere-like HSAT3 (GGAAT)n motif (Fig. 4H) that is commonly seen on Chr20q11 (Altemose et al., 2014). These findings indicate that Chr20q11.21 might be prone to homology-based structural instability at a relatively high frequency, and then the selective advantage conferred by the additional copy of *BLC2L1* drives the expansion of cell lines containing the duplication.

To better understand their potential functional consequences, we next mapped all large structural variants to the genome (Fig. 4I-K). We found that 13/143 (9%) hESC cultures contained aneuploid cells, all but one of which involved chromosomal gain and were predicted to be present at a cellular fraction of 7-66% (Table S3B). Six of these hESC lines showed duplication of Chr12, which has been shown to be recurrent in hPSC cultures (Amps et al., 2011). In addition to aneuploidy, we observed eight fixed large duplications and 20 more present in a fraction of cells, as well as six fixed large deletions and six large deletions present in only a subset of cells (Table S3B). We also found that 14/143 (10%) hESC lines carried CN-LOH events in a subset of cells, of which five involved the entire q arm of chromosome 9 (Fig. 4K). This structural variant has not previously been reported to be recurrent in hPSCs but since we had also previously observed it arising *de novo* upon gene editing (Kiskinis et al., 2014) it is likely to represent a novel, recurrent, culture-acquired structural variant (Fig. 6B).

We also observed the duplication of a sub-region of Chr1q in four cell lines (Fig. 4I), as has been previously reported (McIntire et al., 2016). Since this region has not been finely mapped to the best of our knowledge, we combined our data with an independent set of human induced pluripotent stem cell lines from the HipSci resource (Kilpinen et al., 2017) (data accession: EGAD00010001147), and found 11 more unique cell lines with duplications over this interval. Together, these cell lines defined a minimally duplicated region at Chr1 ~203,408,100 - 204,572,300 (hg19 assembly), corresponding to the cytogenetic location Chr1q32.1 (Fig. 4L). This region contains several candidate genes worthy of future investigation that may confer selective advantage when duplicated, including *KDM5B*, *PIK3C2B*, and *MDM4* (Fig. 6B, Table S3H).

We were surprised to observe two cell lines displaying patterns of mosaic CNV calls consistent with “trisomy rescue” of chromosomes 5 (Genea48) or 16 (HUES71) (Fig. S3H). These trisomies must have resulted from meiotic nondisjunction since they have three distinct haplotypes on segments of these chromosomes, as opposed to two imbalanced haplotypes that might arise from mitotic errors. These findings indicate that at least some structural variants we observed were present in human embryos at the time of hESC derivation. Trisomic cells may be “rescued” to a diploid state by losing one of the excess chromosomes to either restore chromosomal balance or cause uniparental disomy, where both chromosomes are derived from the same parent as seen in Prader Willi Syndrome and Angelman Syndrome. Our findings suggest that meiotic trisomy rescue may also occur *in vitro*, providing an unique opportunity to explore the biology of a process that cannot be readily studied in primary human tissue or hiPSCs, highlighting the enduring relevance of hESCs for human disease modelling. Indeed, meiotic nondisjunction is common in oocytes from older mothers, with chromosome 16 trisomy present approximately 1% of human embryos, typically resulting in miscarriage.

Finally, we wondered whether small (<1 Mbp) CNVs might wholly or partially affect genes of likely functional relevance for hESC biology (Table S3I,J). We found that one cell line (WIBR2) carried a small heterozygous deletion encompassing *TP53* that was present in just 5/234 (2%) of control samples. Furthermore, two unrelated cell lines (CSES6 and CSES25) carried distinct heterozygous deletions encompassing *EP300* that were not observed in human controls (Table S3I). The EP300 gene product acetylates and stabilizes p53 (Gayther et al., 2000), suggesting its reduced dosage could contribute to reduced p53 activity. Overall, our results suggest that small culture-acquired CNVs may functionally impact hPSC biology.

### Frequency of SNVs in hESCs

Missense and loss-of-function (LoF) SNVs can profoundly alter cellular function by affecting both coding regions and functionally important non-coding regions of the genome. Although individual SNVs sufficient to cause human disease are rare in a given individual, in aggregate they affect over 300 million people worldwide (Nguengang Wakap et al., 2019). We therefore predicted that a fraction of hESCs we sequenced would carry SNVs that impact their utility in certain applications. To identify such variants in hESCs, we considered SNV calls from the autosomes and X chromosome that were supported by high confidence and high quality (HC-HQ) sequence data using filters similar to those used by Karczewski and colleagues to analyse 15,708 whole genome sequences in the gnomAD database (Karczewski et al., 2019) (Fig. 5A). After applying HC-HQ filters, we observed an average SNV burden per hESC line of 244 LoF variants and 11483 missense variants, which was indistinguishable with that observed in gnomAD whole genome sequences from humans of diverse ancestries (Fig. 5B).

**Figure 5.**
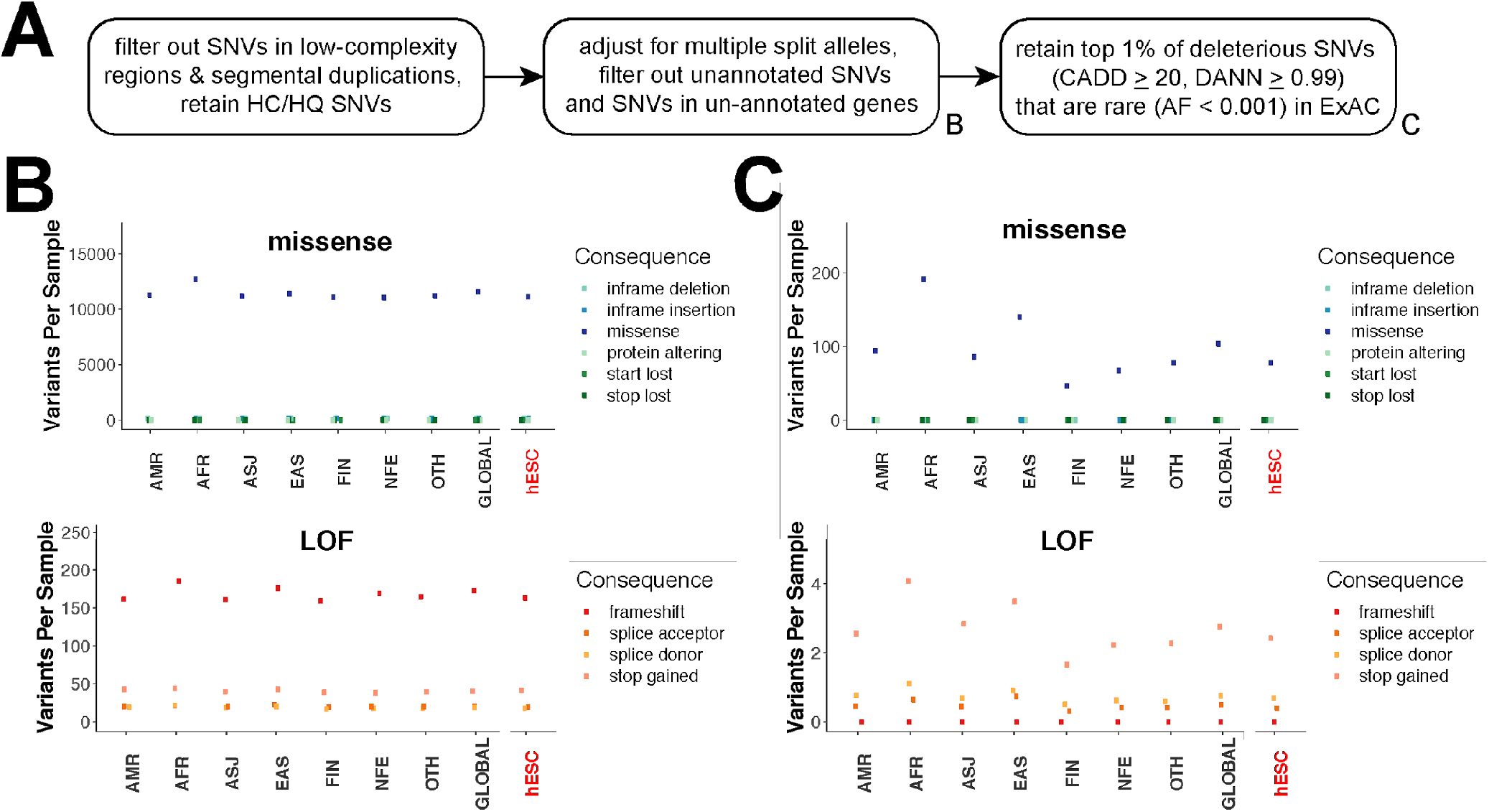
The overall SNV burden in hESCs resembles that of human populations. **A)** Workflow for SNV identification and prioritization based on sequencing quality, bioinformatic prediction of deleteriousness, and ExAC allele frequency. HC/HQ, high-confidence/high-quality. **B)** Per-sample burden of missense and LoF SNVs passing HC/HQ filters across different human ancestries and in the analyzed hESCs. **C)** Per-sample burden of rare and predicted deleterious missense and LoF SNVs. AMR, Ad-mixed American ancestry; AFR, African ancestry; ASJ, Ashkenazi Jewish ancestry; EAS, East Asian ancestry; FIN, Finnish ancestry; NFE, non-Finnish European ancestry; OTH, other ancestry; GLOBAL, all whole genome samples in gnomAD; hESC, hESC samples in this study.

Given that deleterious variants are rare in the general population due to negative selective pressure, we restricted our analysis to variants present at an allele frequency (AF) of less than 0.001 (0.1%) among 60,706 exomes represented in the ExAC database (Lek et al., 2016), which are non-overlapping with the gnomAD whole genomes. To conservatively enrich for likely deleterious variants among these rare SNVs, we used the bioinformatic prediction tools “Combined Annotation Dependent Depletion” (CADD) (Kircher et al., 2014) and “Deleterious Annotation of genetic variants using Neural Networks” (DANN) (Quang et al., 2015) to identify the variants predicted to be among the top 1% most deleterious in the human genome (CADD phred >20 and DANN >0.99). In the hESC lines we analysed, 9,982 SNVs met these criteria (Table S4A). When we compared the burden of these predicted deleterious variants in hESCs to similarly-filtered whole genomes from gnomAD, we again did not observe an enriched burden of deleterious SNVs in hESCs (Fig. 5C). Indeed, the burden of missense and LoF variants per genome for hESCs was consistent with that seen in ancestry-matched (non-Finnish European, NFE) samples. Together, these findings are consistent with the null hypothesis that, under the conditions tested, hESCs do not accumulate an excess burden of SNVs that is detectable above the sampling noise of normal inter-individual genetic variation.

### Cancer-associated SNVs and structural variants

Genetic variants associated with cancer are of particular interest to the stem cell community since they might alter hESC genomic stability and growth characteristics, disrupt hESC differentiation and cellular phenotypes in differentiated cells, or increase the risk of cancerous growths arising from hESC-derived cells after transplant. We therefore asked if any of the SNVs observed in hESCs fell within genes having a documented “Tier 1” activity relevant to cancer as annotated in the Catalogue of Somatic Mutations in Cancer (COSMIC, https://cancer.sanger.ac.uk/cosmic) (Tate et al., 2019). We then asked which of the 382 variants meeting these criteria had been observed in human cancers in COSMIC at least twice (n=51) and were bioinformatically predicted to be cancer-causing by Functional Analysis Through Hidden Markov Models (FATHMM, http://fathmm.biocompute.org.uk/cancer.html) (Shihab et al., 2013). This analysis revealed 14 unique heterozygous missense variants across 10 genes in 15 hESC lines (Table S4B, Fig. 6B), including three of five mutations in *TP53* that we had previously identified by exome sequencing (Merkle et al., 2017). Of the remaining 11/14 SNVs we identified, several suggested the recurrent involvement of the p53 and DNA damage response pathways. In particular, we observed two cell lines (HUES44 and UM77-2) carrying variants in the cell cycle checkpoint kinase ataxia telangiectasia mutated (*ATM*), whose gene product phosphorylates p53 and other downstream targets in response to DNA damage (Maréchal and Zou, 2013). We also observed a likely pathogenic mutation in the *BRCA2* gene in cell line HUES53, whose gene product associates with p53 and helps maintain genomic stability via the homologous recombination pathway for DNA damage repair (Roy et al., 2012). These findings are consistent with our earlier discovery of rare, small CNVs affecting *TP53* and *EP300*.

**Figure 6.**
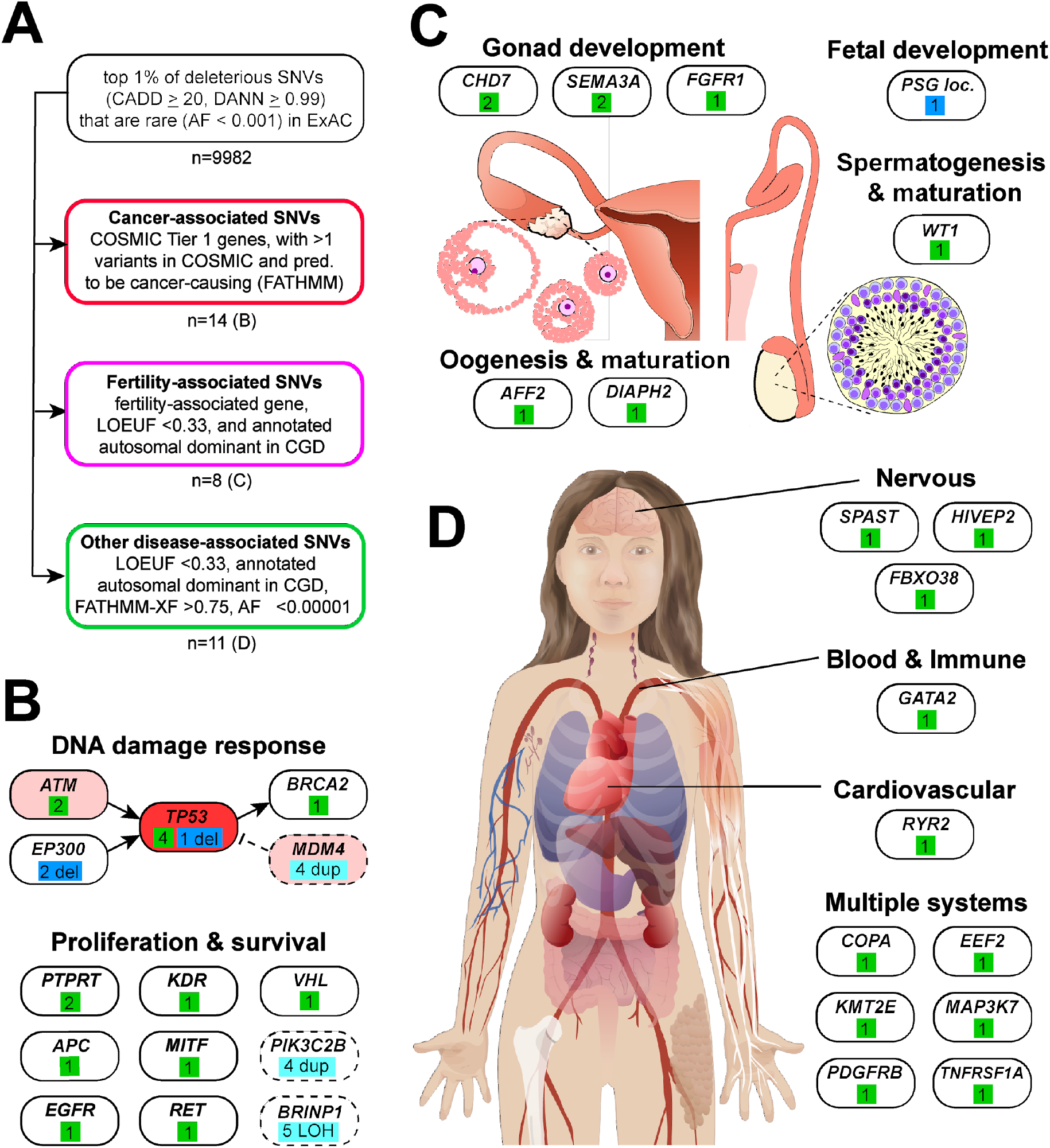
Genes and systems affected by likely deleterious SNVs in hESCs. **A)** Schematic of approach used to identify disease-associated genetic variants in hESCs, and number of variants passing these filters. **B)** Analysis of cancer-associated variants suggests broader involvement of the p53 pathway (KEGG, red shading). Green boxes denote SNVs, and blue boxes denote small (dark blue) or large (light blue) structural variants. Candidate genes implicated by large structural variants are indicated by dashed lines. **C)** Numerous fertility-associated genes carried deleterious mutations, suggesting potential causes of sub-fertility in some of the couples who donated embryos for hESC derivation. **D)** Other SNVs affected genes associated with autosomal dominant disease affecting multiple body systems and cell types generated from hPSCs.

Other hESCs carried likely pathogenic variants in genes relevant to proliferation and apoptosis. HUES42 contained a variant in the von Hippel-Lindau tumor suppressor (*VHL*), whose heterozygous loss of function leads to a dominant cancer predisposition syndrome (Kaelin, 2002). A separate hESC line (CT2) carried a variant in microphthalmia-associated transcription factor (*MITF*). MITF directly regulates the transcription of multiple genes, including the anti-apoptotic gene *BCL2* (McGill et al., 2002) whose paralog *BCL2L1* likely drives the recurrent duplication of Chr20q11.21 in hPSCs (Avery et al., 2013; Nguyen et al., 2014).

In addition, we observed several pathogenic variants in genes associated with cell adhesion and signal transduction. Two cell lines (UM14-1 and CSES25) carried variants in protein tyrosine phosphatase receptor type T (*PTPRT*), a transmembrane protein that associates with members of the cadherin and catenin family of cell adhesion molecules (Besco et al., 2006) and dephosphorylates STAT3 to reduce the expression of its target gene *BCL2L1* (Zhang et al., 2007). We speculate that these *PTPRT* variants may be advantageous to hPSCs by increasing *BCL2L1* expression. Similarly, line UM121-7 carried a variant in adenomatous polyposis coli (*APC*), whose gene product regulates cell proliferation and adhesion via beta-catenin (Aoki and Taketo, 2007), suggesting this cell line might display aberrant responses to WNT signaling; mutations in *APC* are a primary driver of colon adenomas, and this line might therefore by inappropriate for many transplantation applications (but useful for studies of precancerous states). Cell line WA21 carried a pathogenic variant in the protooncogene *RET*, which encodes a receptor tyrosine kinase in which gain-of-function variants promote endocrine tumors and other cancer types (Mulligan et al., 1993), while cell lines CT3 and a line from UCLA carried mutations in the receptors for epidermal growth factor (EGF) and vascular epithelial growth factor (VEGF), respectively.

Finally, we asked if the COSMIC-associated variants we identified in hESCs were included among the 23 genes recurrently mutated in hESCs over time in culture (Avior et al., 2019). We identified variants in 9 of these genes, including *FAT1* (n=6), *TP53* (n=4), *APC* (n=3), *ARID1A* (n=3), *EGFR* (n=1), *MYH9* (n=1), *PCM1* (n=1), *RNF213* (n=1) and *VHL* (n=1). Notably, the *RNF213* variant was present in a homozygous state in line WIBR6. It will be interesting to see in future studies whether mutations in these genes alter hPSC growth or survival (Table S4B).

### Fertility-associated variants

The majority of hESCs we studied here were derived from donated embryos there were excess to the needs of couples seeking assisted reproduction at *in vitro* fertilization (IVF) clinics. We thus considered whether genetic variants affecting fertility would be present among the sequenced hESC lines. It is of practical importance to identify such cell lines, since there is considerable interest in differentiation of hPSCs into germ cells for studies of human meiosis and gametogenesis (Sasaki et al., 2015; Zhou et al., 2016). To identify such infertility-associated genetic variants in hESCs, we examined the intersection between a list of genes associated with infertility from the literature (Mallepaly et al., 2017; O’Flynn O’Brien et al., 2010; Venkatesh et al., 2014), and genes in which inactivating heterozygous mutations are under strong negative selective pressure and are therefore likely to cause dominant disease as identified using the loss-of-function observed/expected upper bound fraction (LOEUF) metric from studies of human genome variation (Karczewski et al., 2019). We conservatively set LOEUF to 0.33 or less, revealing variants in 12 fertility-associated genes meeting these criteria. To increase our confidence in the disease relevance of these variants, we further restricted our analysis to genes whose disruption is associated with autosomal dominantly-inherited disease in the manually-curated Clinical Genomic Database (CGD, https://research.nhgri.nih.gov/CGD/) and identified 8 variants affecting 6 of the 12 previously identified genes (Fig. 6C, Table S4D). These included a variant in Wilms Tumor 1 (*WT1)*, whose dysfunction is associated with abnormal sperm production and female subfertility (Nathan et al., 2017; Niederberger, 2014). We also identified two independent missense variants in Chromodomain Helicase DNA Binding Protein 7 (*CHD7*) whose disruption can cause Kallmann syndrome (hypogonadotropic hypogonadism), which is characterized by underdeveloped testes or ovaries due to defects in cell populations in the hypothalamus or pituitary gland that regulate sexual development (Jongmans et al., 2009). We further observed three variants in genes associated with autosomal dominant hypogonadotropic hypogonadism; two affecting *SEMA3A* (Hanchate et al., 2012) and a likely inactivating variant in *FGFR1* (Dodé et al., 2003). Both *SEMA3A* variants are strongly predicted to be pathogenic by CADD and DANN (scores >30 and >0.999, respectively) as well as FATHMM-XF (score >0.7) (Rogers et al., 2018), and one of these variants affected each of the 3 sibling lines in a trio we identified (Fig. 2D), raising the intriguing possibility that it might explain the cause of sub-fertility in the couple who donated their embryos to generate these hESC lines.

We also observed two cell lines (HUES62 and KCL020) with variants in the X chromosomal genes *DIAPH2* and *AFF2*, respectively. The disruption of these genes is associated with premature ovarian failure, as well as other conditions including mental retardation. Groups seeking to differentiate hESCs into oocytes may wish to avoid these cell lines. We also identified a cell line (WA26) that carried a homozygous deletion of the pregnancy-specific glycoprotein (PSG) gene cluster. PSG genes are still poorly understood but encode the most abundant fetal proteins found in the maternal bloodstream and likely have immunomodulatory functions important for fetal growth and survival (Lisboa et al., 2011; Martinez et al., 2013).

### Disease-associated SNVs

Finally, we screened for variants that might cause disease-relevant phenotypes *in vitro* or compromise the safety of hESC-based regenerative medicine. We focused on genes predicted from human-genetic variation data to be intolerant to loss-of-function (LOEUF <0.33, n=2000) and that had been annotated as associated with autosomal dominant human disease in CGD (n=324). To restrict ourselves to the variants in autosomal genes most likely to be pathogenic, we considered those bioinformatically predicted to be pathogenic by FATHMM-XF (score >0.75, n=146) (Rogers et al., 2018) and DANN (score >0.999, n=36). Finally, we further filtered these variants to those that were extremely rare in ExAC (AF <1×10^−5^) in order to enrich for those variants that were damaging enough to be subject to purifying selection. These filters revealed 11 variants whose disruption is associated with disease *in vivo* and might also affect cell types commonly generated from hPSCs *in vitro* (Fig. 6D, Table S4E). We found that some variants affected genes required for normal development and might therefore interfere with the generation of specific cell types. In particular, one cell line (KCL019) carried a variant in *GATA2*, a transcription factor involved in immune cell development associated with immunodeficiency and leukemia when disrupted (Collin et al., 2015). Groups seeking to generate hPSC-derived blood cells (Moreau et al., 2016; Sugimura et al., 2017) may wish to avoid cell lines carrying this or similar variants.

Other variants likely affect the function of specific cell types. KCL032 carried a mutation the gene encoding the ryanodine receptor RYR2, which is enriched in the heart where it mediates the release of intracellular calcium from sarcomeres during cardiomyocyte contraction. Mutations in this gene have been associated with autosomal dominant cardiac arrhythmia and tachycardia (Priori and Napolitano, 2005), suggesting that cardiomyocytes differentiated from hPSCs carrying this variant might display abnormal function *in vitro*. Furthermore, multiple cell lines carried variants that might alter neuronal function. WA19 carried a variant in the spastin (*SPAST*) gene, whose gene product interacts with and severs microtubules, and whose dysfunction can cause the autosomal dominant neurological disease spastic paraplegia characterized by axonal defects (Solowska and Baas, 2015). Similarly, cell line KCL038 carried a variant in *FBXO38*, which encodes a co-activator of KLF7, which has been associated with motor neuron degeneration (Sumner et al., 2013) and HUES69 carried a variant in the lysine methyltransferase *KMT2E* (*MLL5*), which plays a role in regulating gene expression, cell-cycle progression, and genomic stability as well as having been associated with epilepsy and neurodevelopmental defects (Zhang et al., 2017a). A fourth cell line (KCL022) carried a missense variant in *HIVEP2*, whose loss of function is associated with autosomal dominant mental retardation (Srivastava et al., 2016).

We also found variants in genes involved in broader cell biological processes that might cause numerous phenotypes *in vitro* when disrupted. Cell line HUES45 carried a variant in *TNFRSF1A* (*TNFR1*), which encodes a receptor for tumor necrosis factor (TNF) and plays a role in cell survival, apoptosis, and inflammation (Ting and Bertrand, 2016). Interestingly, a second cell line (WA24) carried a missense variant in the MAP kinase family member *MAP3K7* (MEKK1), whose gene product functions downstream of TNFRSF1A and MEK, and that binds to and stabilizes p53 (Chipps et al., 2015). Heterozygous mutations in *TNFRSF1A* or *MAP3K7* have been associated with pathological inflammatory responses or defects in skeletal and craniofacial development, respectively. *In vitro*, these variants may lead to altered TNF-alpha signal transduction and apoptosis. We also observed a cell line (from UCLA) carrying a variant in *COPA*, whose gene product is required for normal retrograde membrane trafficking from the Golgi to the endoplasmic reticulum (ER). COPA disruption causes ER stress and is associated with autoimmune joint, lung, and kidney disease *in vivo* (Watkin et al., 2015) and may therefore cause cellular phenotypes *in vitro* in cells sensitive to ER stress or affect immune cell generation. Furthermore, cell line CHB5 carried a heterozygous missense variant in the gene encoding the eukaryotic translation elongation factor EEF2, which is required for normal rates of protein synthesis (Kaul et al., 2011). Heterozygous mutations in this gene have been associated with spinocerebellar ataxia, and hESCs and their progeny carrying this variant may display altered protein metabolism, with unclear downstream consequences. Finally, we observed a cell line harbouring a variant in the platelet derived growth factor receptor *PDGFRB*. Mutations in this gene have been linked to multiple diseases of abnormal growth, development, and aging (Andrae et al., 2008).

We note that while the discovery of potentially deleterious SNVs is concerning and their potential effects should be carefully considered, many of the specific alleles we found lack definitive experimental evidence demonstrating that they are sufficient to cause disease. Conversely, our stringent bioinformatic selection criteria clearly exclude some functionally relevant variants. For example, filtering on LOEUF excluded all variants in *TP53* and a D90A variant in *SOD1* associated with incompletely penetrant amyotrophic lateral sclerosis (Al-Chalabi et al., 1998) that may be relevant to groups modelling neurodegenerative disease. Furthermore, our sequencing quality filters removed two of the five variants previously identified in *TP53* since they were present at low allelic faction. We therefore provide a full list of likely deleterious SNVs (Table S4A) and developed a web resource to enable groups to mine these data and rationally select cell lines.

### Tools for rational hESC line selection

The breadth of findings that can be garnered from whole genome sequencing data raised the question of how genomic information can best be harnessed by stem cell biologists to rationally select an appropriate cell line for a particular application. Which variants are likely benign, and which might limit the utility of a cell line in a given application? To help the community address these questions, we generated three complementary resources. First, we summarize some of the most relevant results presented in this study, in the form of a convenient lookup table (Fig. 7A). Second, we provide an annotated table of the variants we identified (Tables S1-S4) and deposited the raw sequencing data into controlled access databases (dbGaP, EGA). Finally, we have created a user-friendly online data portal (https://portals.broadinstitute.org/hscgp) (Fig. 7B, Movie S1) that enables users without computational expertise to readily search for sequence variants of interest among sequenced hESC lines. For example, a search for *TP53* reveals all variants in the gene that were detected in the sequenced cell lines, the names of those cell lines, as well as bioinformatic predictions about the likely consequences of these variants. Search results can be exported for further analysis. Specific cell lines can also be interrogated for the presence of variants of interest, and raw sequencing alignments can be visualized via the integrative genomics viewer IGV (Robinson et al., 2011).

**Figure 7.**
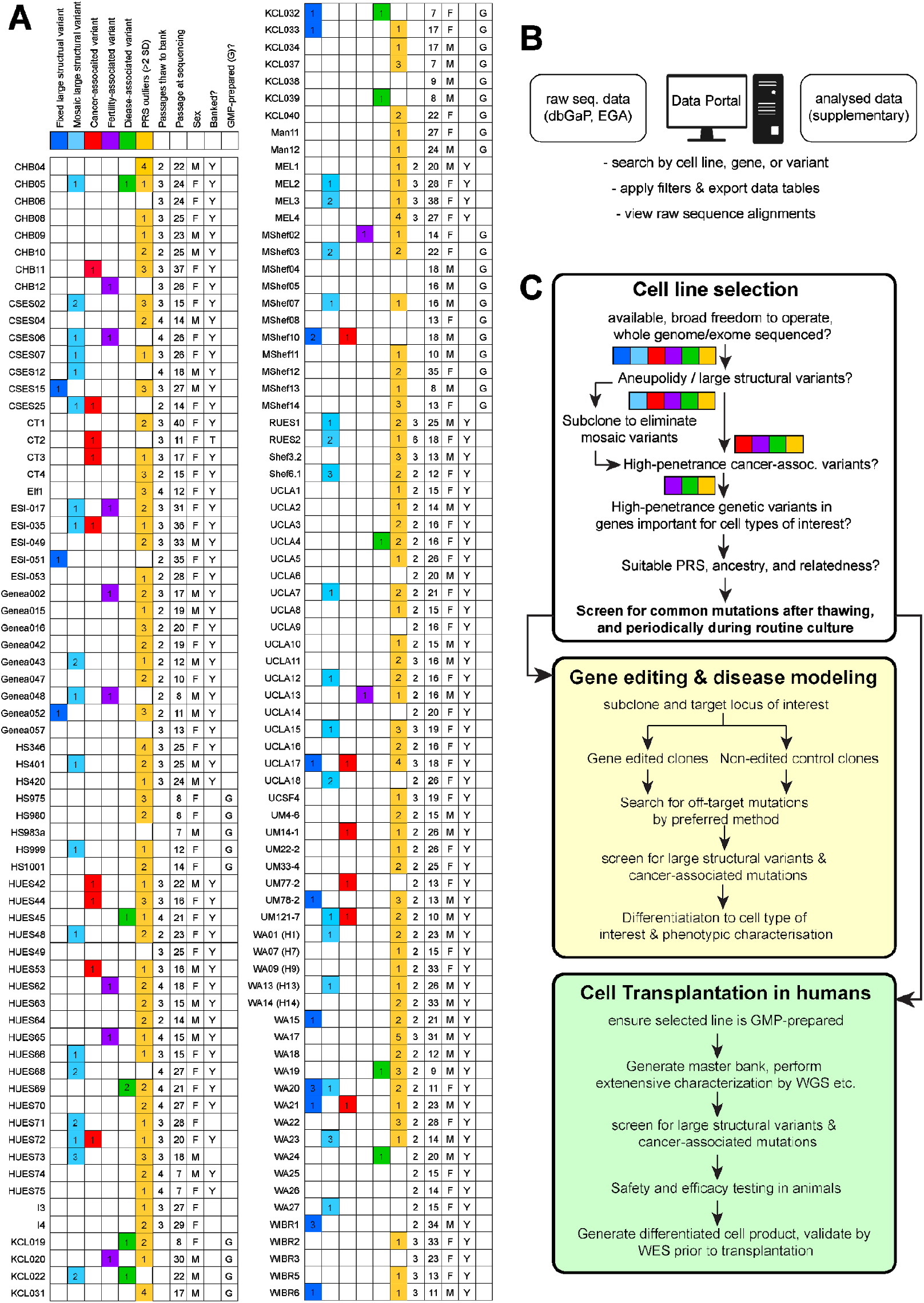
Genetically-informed rational cell line selection. **A)** Graphical summary of the number of large fixed and mosaic structural variants (dark and light blue, respectively), SNVs likely associated with cancer (red), fertility (magenta) or other diseases (green), “outlier” PRS, and summary data for each analyzed hESC line. **B)** As a resource for the field, raw sequencing data are deposited in controlled-access databases, analyzed data are provided as supplementary data files associated with this study, and user-friendly data portal enables non-specialist groups to explore and extract data of interest, down to the level of the primary sequence traces. **C)** Suggested workflow for rational hPSC line selection and monitoring of genomic integrity for research and clinical applications.

## DISCUSSION

Techniques commonly used to evaluate the genetic integrity of hESCs, including high-density SNP microarrays and karyotyping by G-banding, have limited spatial resolution and limited power to detect mosaic events (Baker et al., 2016). Here we show that high-coverage (> 25x) WGS enables the robust identification of potentially relevant structural and single nucleotide variants. As the price of whole genome sequencing is steadily dropping and provides considerably more information on genetic variation than many other methods, our experience suggests that WGS may gradually become the tool of choice for genetic analysis of hPSCs.

While we strove to identify the most relevant genetic variants present in hESC lines, our analyses were not exhaustive. We did not consider inversions, translocations, repetitive genomic regions, mitochondrial DNA sequences, epigenetic differences, or variants on the Y chromosome. While the WGS data used in this study provides an unprecedented view of stem cell genomes, its short read length (150 base paired-end reads) coupled with the inherent variation in sequencing read depth due to DNA replication (Koren et al., 2014) does have some limitations that might be mitigated in part by future studies using long-read sequencing technologies. For instance, our analysis was currently limited to variants <~50 bp or >~1.1 kbp, and our analysis structural variant mosaicism was limited to variants >~1 Mbp. In addition, the accuracy of HLA haplotype estimation is constrained by the limited number of informative SNPs, and haplotypes should be verified prior to use in any downstream application (De Bakker et al., 2006). We therefore encourage groups to interrogate the full dataset for relevant variants in their cell line(s) of interest using the provided web resource (https://portals.broadinstitute.org/hscgp) or by re-analysis of the raw data (dgGaP, EGA). We also anticipate that groups may wish to utilise the data sets presented here in combination with transcriptomic data to map expression quantitative trait loci (eQTL) in hESCs or their differentiated progeny.

### Genetic background and relatedness

We found that all but a few of the hESC lines we sequenced were of European ancestry, highlighting the need to expand the diversity of stem cells lines available in order to facilitate study of genetic variants found in individuals of distinct ancestries. We were also surprised that 33% of sequenced hESC lines were siblings. While these related cell lines should not be treated as independent controls in biological experiments, they may be valuable for examining the impact of genetic inheritance on transcriptional and phenotypic outcomes. Each cell line also carried a distinct PRS profile based on combinations of thousands of common genetic variants. The predictive value of PRS score calculations will likely increase as population sequencing studies include larger numbers of more deeply-phenotyped individuals. Selecting lines based on their PRS may improve case-control and isogenic disease models, since hPSC genetic background drives transcriptional variability (Kilpinen et al., 2017; Rouhani et al., 2014b), and incompletely penetrant phenotypes may only be detectable on a genetically predisposed background, as is well-known in animal models (Ayadi et al., 2012).

### The origin and impact of structural genetic variants

We found that almost a third of hESC lines in this study carried large structural variants, of which approximately half were mosaic (Fig. 4J,K, Fig. 7A, Table S3B). Mosaic variants might arise in culture and confer selective advantage to affected cells, leading to the expansion and eventual fixation of the variant. Indeed, of the nine hESC lines carrying advantageous duplications at Chr20q11.21, six were mosaic and three were fixed (Fig. 4B). Alternatively, mosaicism could arise from negative selective pressures, since approximately half of all preimplantation human embryos carry large structural genetic variants (Fragouli et al., 2019) and we observed two instances of apparent trisomy rescue (Fig. S3H). Fixed variants might have arisen in culture and reached fixation or might be inherited as seen in the sibling cell lines KCL032 and KCL033 that carried identical large duplications on Chr5 (Table S3B).

Our studies confirmed the recurrence of duplications on Chr1, Chr12, and Chr20q11.21 (Fig. 4J). We found that Chr20q11.21 duplication breakpoints contain a common centromere-like HSAT3 motif, suggesting that this chromosomal region is predisposed to microhomology-based copy number changes (Fig. 4H,I). We also found that four hESC and 11 hiPSC lines carried a minimally duplicated ~1.2 Mbp region on Chr1q32.1 containing several genes that might confer selective advantage, either alone or in concert (Fig. 4M). These genes include *MDM4* which is frequently overexpressed in cancer and whose gene product directly binds to and inhibits p53 (Fig. 6B) (Francoz et al., 2006), the PI3 kinase *PIK3C2B* which is frequently duplicated in malignant breast cancer and regulates cell proliferation and DNA damage response (Zhang et al., 2017b).

We also discovered a novel, recurrent CN-LOH event on Chr9q (Fig. 4L) that would alter the expression levels of imprinted or differentially methylated genes. Indeed, Chr9q LOH is seen in most bladder cancer cases (Hirao et al., 2005), and sometimes also observed in non-cancerous esophageal mucosa samples (Yizhak et al., 2019) and in tissue adjacent to head and neck squamous cell carcinoma (Jakubek et al., 2020), suggesting a role cancer formation. Together these studies suggest Chr9q may contain genes expressed in an allelically unbalanced manner that affect cell growth or survivalFor example, *BRINP1*, also known as *DBCCR1* and *DBC1* (deleted in bladder cancer 1) located at Chr9q33.1 is frequently deleted or hypermethylated in cancer (Izumi et al., 2005) (Fig. 6B), as is the microRNA miR-181a2/181b2 at Chr9q33.3 (Fig. 6E) which suppresses cell cycle progression likely by targeting the PI3K/AKT/MTOR pathway (Mei et al., 2017). Future transcriptional and epigenetic profiling studies of hPSCs may reveal specific genetic variants associated with the recurrence of this variant.

### Disease-associated SNVs and suggestions for rational hESC selection

We sought to identify sequence variants likely to affect the function of hESCs or their derivatives using combination of gene-level and variant-level filters based on manually-curated databases and bioinformatic prediction algorithms (Fig. 6B-D). This approach avoids potential selection bias but we note that databases are not comprehensive and prediction algorithms are imperfect, and it is also difficult to predict which phenotypes will result from specific genotypes. Indeed, there are no hard rules governing which mutations might disqualify a cell line for a given purpose and polyploid cells have been knowingly transplanted into humans without apparent ill effects (Nelson et al., 2002). While we feel that most groups would agree that a cell line carrying a *TP53* mutation conferring a lifetime cancer risk of nearly 100% (Malkin, 1993) should not be transplanted into patients if a suitable alternative exists, other variants should be considered on a case-by-case basis depending on research needs. In contrast, damaging variants associated with fertility or other disease (Fig. 6B-C, Table S4) will likely be of interest and perhaps even value to groups studying gametogenesis or modelling specific diseases *in vitro*.

To facilitate the rational selection of hPSC lines based on genetic criteria, we therefore suggest a scheme (Fig. 7C) that can be adapted to achieve the appropriate balance of risk and benefit for a particular application. We illustrate decision points that many groups working with hPSCs will likely encounter (Fig. 7C). First, we reason that most groups will prefer to work with cell lines having minimal restrictions on the freedom to use or share the lines and their derivatives, and whose genomic structure has been extensively characterized. Second, since aneuploidy or rare, large structural variants would affect the expression of many genes with unpredictable consequences, we suggest avoiding such lines for both research and clinical use, although it might be possible “rescue” lines with mosaic variants by sub-cloning or by requesting an earlier passage line from the provider that may lack the variant. Third, most groups would avoid cell lines carrying cancer- and disease-associated variants whose predicted effect size exceeds pre-established criteria, which will vary by application. Notably, some groups may wish to take advantage of genetic disease predisposition rather than using this as a criterion for excluding cell lines. Fourth, the PRS for a trait of interest, ancestry, and relatedness of the cell line may reveal cell lines are most likely to display desired traits.

### Recommendations for monitoring hPSC genomes

We found that both structural and sequence variants likely to compromise hPSC function are widespread, and some were likely acquired in culture. This fact highlights a pressing need to reduce the selective pressures that apparently exist in commonly used culture conditions. A clear pattern among many of these mutations was their effects upon TP53, EP300, and other genes involved in response to DNA damage. These findings underscore the importance of regularly monitoring the genomic integrity of cultured hPSCs. The criteria for monitoring genomic integrity might be more stringent for clinical applications, but we recommend that groups performing basic research could streamline tissue culture workflows and achieve more reproducible results by following the steps outlined in Figure 7C. Briefly, since hPCSs acquire genetic variants over time in culture, we suggest selecting and subcloning a deeply characterized cell line with suitable genetic properties, confirming the absence of undesired genetic variants, and then expanding one subclone to generate a large “master bank” of frozen vials of known genetic status. After confirming that potentially deleterious mutations did not arise during cell line expansion, “working stock” vials could be periodically drawn from the bank and then discarded within 5 passages after thawing to minimize genetic drift and reduce the need for regular sequencing. If lines need to be cultured for longer periods, we recommend that they be routinely sequenced every 5-10 passages to test for the emergence of potentially deleterious mutations and to safeguard against accidental cell line cross-contamination (Horbach and Halffman, 2017). We also suggest performing genomic analysis after gene editing and other manipulations that force cells through clonal bottlenecks.

Together, our analysis demonstrates that the overall numbers of small CNVs and SNVs identified in hESCs are indistinguishable from those seen in somatic cells from a similarly scaled population of donors, validating hESCs as a powerful tool to study human development and disease and as a useful source of clinically important cell populations. Despite this global similarity, we uncovered a wealth of genetic variants unique to each cell line, including large structural variants and potentially deleterious SNVs in disease-associated genes that might compromise the utility of certain lines for certain applications. We anticipate that the data provided here will become increasingly valuable as our understanding of genotype-phenotype relationships steadily improves. By providing a searchable online data portal enabling individuals with any levels of computational expertise to make use of the resource we report here, we hope that the reproducibility of research findings from hESC studies and their ultimate use in clinical applications will be improved.

## AUTHOR CONTRIBUTIONS

FTM, SG, SM, and KE conceived of the project and wrote the manuscript with contributions from GG, BH, and SK. FTM led efforts to acquire and sequence hESCs and to analyze and interpret sequencing data together with SG, who coordinated bioinformatic analysis and the development of web resources. GG performed ancestry, relatedness, and CN-LOH analysis. CD and GG performed PRS analysis. FTM and GG identified HLA haplotypes and disease-relevant genes present in clinical databases. KK, GG, and SG performed LoF analysis and overall SNV burden under the guidance of DM. BH, SK, and GG performed structural variant analysis. SG developed SNV analysis pipelines and worked with FTM to develop criteria for variant prioritization. CP and MP generously provided access to unpublished WGS data from individuals of Latin American and African American ancestry unaffected by schizophrenia for CNV analysis.

## ACKNOWLEDGMENTS

We are grateful to the many institutions from around the world who generously provided their cell lines and supported the publication of the results, including Children’s Hospital Corporation, Cedars-Sinai Medical Center, University of Connecticut, BioTime Inc., Genea Biocells, Karolinska Institute, Harvard University, Technion R&D Foundation, King’s College London, University of Manchester, University of Queensland, University of Sheffield, The Rockefeller University, University of California Los Angeles, University of California San Francisco, University of Michigan, WiCell Research Institute, and the Whitehead Institute for Biomedical Research. We thank Diane Santos, Melissa Smith, Kristen Elwell, Mary Anna Yram, Stacey Ellender, Liz Bevilacqua, Diane Gage, and Anna Neumann for their assistance with acquiring hESC lines, coordinating the sequencing and genotyping workflows, and for preparing data for submission to data repositories. The Genomics Platform at the Broad Institute of MIT and Harvard performed sample preparation, sequencing, and data storage. Web tools were developed with the help of the KDUX Data Sciences Platform at the Broad Institute of MIT and Harvard. Costs associated with acquiring and sequencing hESC lines were supported by the Howard Hughes Medical Institute and the Stanley Center for Psychiatric Research at the Broad Institute of Harvard and MIT. AFS, SM, and KE were supported by grants from the NIH [HL109525], [5P01GM099117]. FTM is a New York Stem Cell Foundation - Robertson Investigator, and this research was supported by The New York Stem Cell Foundation [NYSCF-R-156], the NIH [5K99NS083713], the Wellcome Trust and Royal Society [211221/Z/18/Z], and the Chan Zuckerberg Initiative [191942].

## MATERIALS AND METHODS

### Whole genome sequencing and genotyping

hESCs were acquired and cultured as previously described (Merkle et al., 2017). Briefly, cell pellets of approximately 1-5 million cells digested overnight at 50C in 500 μl lysis buffer containing 100 μg/ml proteinase K (Roche), 10 mM Tris pH 8.0, 200 mM NaCl, 5% w/v SDS, 10 mM EDTA, followed by Phenol:Chloroform precipitation, ethanol washes, and resuspension in 10 mM Tris buffer (pH 8.0). Genomic DNA was then transferred to the Genomics Platform at the Broad Institute of MIT and Harvard for Illumina Nextera library preparation and sequencing on the Illumina HiSeqX platform with paired-end 150 base reads. Sequencing reads were aligned to the hg19 reference genome using the BWA MEM alignment program. Genotypes from WGS data were computed using best practices from GATK software (McKenna et al., 2010) version 3.1. Data from each cell line was independently processed with the HaplotypeCaller walker and further aggregated with the CombineGVCFs and GenotypeGVCFs walkers. Genotyped sites were filtered using the ApplyRecalibration walker.

### Determination of ancestry, genetic relatedness, and HLA haplotype diversity

Genotypes from the cell lines at sites in common with sites genotyped in the 1000 Genomes Project Phase 1 (Consortium et al., 2012) and with a minor allele frequency (MA) of at least 1% were selected for relatedness and ancestry analysis. For relatedness analysis, selected sites were pruned using PLINK software (Chang et al., 2015) (“--indep 50 5 2” command option) and estimates for amount of IBD1 and IBD2 regions were computed (“--genome gz” command option). Sample pairs were considered directly related (parent-child or full sibling) when estimates were between 35% and 70%.

For ancestry analysis, selected sites were extracted from the dataset, merged with 1000 Genomes Project genotypes, and pruned using PLINK2 software (“--indep 50 5 2” command option). Principal component analysis was then performed for this combined dataset by computing the pairwise relationship matrix across all subjects (using the plink command “--make-grm-bin”), and computing the principal components using GCTA software (Yang et al., 2011). Global ancestral components for European, African, Native American, and Asian ancestry were estimated from the first three principal components using a linear model trained by assigning full European, African, and Asian ancestry to the appropriate 1000 Genomes population samples, and assigning estimated ancestry proportions to Latino samples using available published estimates (Consortium et al., 2012; Maples et al., 2013).Human Leukocyte Antigen (HLA) genotype was ascertained by genotyping hESCs for SNPs associated with HLA haplotypes in the CEU ethnic group (De Bakker et al., 2006).

### Polygenic risk score (PRS) computations

To compute PRSs, we used summary statistics from studies performed on individuals predominantly of European ancestry. For each phenotype with available summary statistics, we computed the score for all available markers with association p-values in the original study (Bulik-Sullivan et al., 2015a) (less than 0.5 (e.g. 1,218,732 for BMI). PRS were computed with established methodologies (Purcell et al., 2009) and PLINK software (Chang et al., 2015) using genotypes from WGS data. To control for potential methodological biases, we also calculated PRSs using high-coverage WGS data from unrelated human samples including 383 schizophrenia (SCZ) cases and 489 SCZ controls (ripke 2014) and adjusted raw PRSs by regressing the first 12 principal component values to correct for slight biases introduced in the computation of the summary statistics. Principal components beyond the first 12 did not show significant correlation with raw PRS. Raw PRS values were normalized such that the distribution of PRS scores from SCZ control subjects had zero mean and unitary standard deviation (SD).

### CNV calling from read depth variation in WGS data

Copy number variants were ascertained and genotyped using Genome STRiP (Handsaker et al., 2015) version 2.0 (r2.00.1587) (http://www.broadinstitute.org/software/genomestrip/), using the CNVDiscoveryPipeline with default settings. This software examines average depth of coverage (DOC) genome-wide to identify chromosomes and genomic regions whose normalized DOC deviated from the expected copy number of 2 for autosomes, and 2 or 1 for the X and Y chromosomes. After initial CNV calling, quality was assessed using signal intensity data from Omni 2.5 Microarrays run on the 500 individuals from a control cohort. Using the IRS method (Handsaker et al., 2015) we established separate length thresholds for deletions (length threshold = 1117) and duplications (length threshold = 2750) that achieved a false discovery rate under 3% for both categories of CNV (Figure S3A). By comparing the number of CNVs called in the cell lines to CNVs called in the control cohort, stratified by CNV type, length and ancestry, we estimate that the false discovery rate in the cell lines should be comparable to the estimated false discovery rate of 3% in the control cohort.

To increase CNV calling power and to enhance quality control, CNV analysis was performed on a combined cohort of 130 sequenced hESC lines and a control cohort consisting of 243 human samples from primary blood or lymphoblastoid cell lines (LCL) that had undergone WGS on the same platform (Illumina HiSeqX) and to similar depth as the hESC lines (Pato et al., 2013). We then separated CNVs into small (1117 bp - 1 Mbp) and large (> 1 Mbp) categories since large variants were uncommon among similarly-sequenced human samples and therefore more likely to be culture-acquired and potentially deleterious, and manually confirmed all large CNV calls from hESCs and human control samples.

### Detection of large fixed and mosaic structural variants

To detect large-scale copy number alterations (> 1 Mbp) by sequencing depth of coverage (DOC), we scanned the genome for segments where one sample had was enriched or depleted in depth of sequencing coverage compared to the other cell line samples. We divided the genome into non-overlapping 100 kbp bins of uniquely-alignable sequence (based on 101bp k-mers) and computed the normalized depth of coverage using Genome STRiP (Handsaker et al., 2015). For each contiguous range of bins, a Z-score was computed for each sample over the genomic interval as:

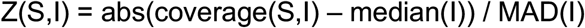

where coverage(S,I) is the normalized depth of coverage for sample S over the interval, median(I) is the median coverage over all samples and MAD is the median-absolute-deviation. We performed a heuristic search to identify candidate CNV intervals with a Z-score > 3 that were local maxima for sample S. We required that candidate CNV intervals have sharp boundaries, such that the depth of coverage for sample S in the next adjacent bin (on both ends of the interval) was more than halfway from the coverage for S inside the interval to the median coverage of the adjacent bin outside the interval. The final set of large CNVs reported were selected to span at least 10 bins (minimum length 1 Mbp) and have a Z-score > 5. The boundaries of each CNV were manually reviewed and adjusted as needed near centromeres or telomeres or to merge adjacent calls. Contiguous but compound variants were considered as one unique event.

To determine if CNVs detected in hESCs were present in all sequenced cells or in a subset of them, we calculated the likelihood that the divergence from an integer copy number could have arisen by chance. We set the P value threshold for detecting mosaicism at 1 × 10^−3^ since human blood samples had only 2/243 (< 1%) samples with smaller P values (Fig. S3B), and otherwise classified CNVs as fixed. CNV calls made from sequencing depth of coverage analysis were supported by allelic coverage at heterozygous sites. This was measured at all heterozygous SNPs observed at least five times across the whole dataset and falling outside of regions exhibiting excessive heterozygosity in the 1000 Genomes project dataset, as these regions might be more prone to mis-mapping. Each remaining heterozygous allele was then phased using Eagle (Loh et al., 2016). Phased genotypes and allelic coverages were then analyzed for allelic imbalances with MoChA (https://github.com/freeseek/mocha). Highly confident calls (LOD>20) of large structural variations (> 1 Mbp) were reported (Table S3). To distinguish acquired CN-LOH events from fixed runs of homozygosity arising from inbreeding or complex chromosomal rearrangement, we considered only events that were mosaic and extended to the telomere, consistent with CN-LOH formation in culture by a single mitotic crossover event.

### Comparison of structural variant calls by SNP array and WGS

All structural variant calls made by SNP array in a published dataset (Canham et al., 2015) were compared to those made in this study that were 500 kbp or larger. Calls made by SNP array had been mapped to the hg38 reference genome, so variant calls were converted to hg19 coordinates to enable comparison to WGS data. We considered the 20 variants detected by Canham and colleagues regardless of their size, and found that 15 of them were confirmed by WGS data. Manual inspection of the five calls made by SNP array only showed that three did not meet our criteria for real CNVs, and two were likely explained by the divergent culture histories of cell lines that were sequenced at different passages.

### Detection of small CNVs and the genes they affect

Regions containing large fixed or mosaic CNVs were excluded from small-scale CNV analysis in affected samples to reduce the potential “fragmentation” of large CNVs into smaller calls. Singleton events were defined as being present only once in any of the other hESC lines, human samples, or in previously described databases such as the 1000 Genomes Project (Auton et al., 2015; Consortium et al., 2012).

To ask whether hESCs might be enriched for specific smaller CNVs, we examined the genome-wide distribution of CNVs < 1 Mbp in hESCs and removed all but one cell line from the sib-ships we identified to control for CNVs shared between first-degree relatives. We measured the frequency of each CNV found in the hESCs in two control cohorts: 212 human samples of European ancestry and 250 human samples of Latino or African American ancestry. We identified events that were present in at least 5 hESC lines but that were absent from the European control samples and found in less than 10% of the other control samples. This analysis identified the known region on Chr20q11.21 as well as four other potentially differentiated CNVs. Manual review of these four additional regions revealed that they are all in small regions of extreme GC-content with excessive sequencing coverage and not likely to be real CNVs.

For each cell line, genes overlapping with 5997 CNVs were identified and annotated using the GeneOverlap module in GenomeStrip 2.0 and GENCODE. The region of overlap (coding vs. exonic vs. intronic) and the type of CNV (deletion or duplication) were also recorded. 2185 CNVs contained at least some part of an annotated gene (Table S3). Limiting CNV overlaps to coding regions and UTRs reduced these numbers to 357 deleted (CN < 2) and 321 duplicated (CN > 2) genes, respectively.

### Identification of loss of function (LoF) and missense SNVs in hESCs

To ensure reliable assignment of variant function and to confidently identify potentially high impact variants, genotypes were called jointly in all hESC lines and genotype data were annotated using Variant Effect Predictor (VEP version 85) in Hail (https://hail.is) with the Loss-of-function Transcript Effect Estimator plugin (LOFTEE, https://github.com/konradjk/loftee) (McLaren et al., 2016). Candidate variants called by VEP were advanced for further analysis and false positive calls were removed from the VEP variant call set by applying similar filters used by gnomAD to identify “high confidence and high quality” (HC-HQ) variant calls (Karczewski et al., 2019). Specifically, we split multiallelic sites and discarded all calls without a “PASS” filter tag as determined by the Variant Quality Score Recalibration tool (GATK). We also excluded variants sites where not a single sample had high genotyping quality as defined by (Depth of Coverage >= 10, Genotype Quality >= 20 and Minor allele fraction >= 0.2). Variants in low complexity regions and segmental duplications were filtered out and only variants that met the gnomAD “PASS” filter criteria or were missing were retained. These filtering steps resulted in a total of 15.5M high quality variants corresponding to approximately 70% of all variants in the data.

Since VEP annotates individual transcripts, only variants on canonical transcripts as defined by Gencode/Ensembl were included in downstream analyses. Variants were binned into synonymous, missense and lofs using the following criteria:

<u>LoFs</u>: “frameshift_variant”, “splice_acceptor_variant”, “splice_donor_variant”, “stop_gained”

<u>Missense</u>: “inframe_deletion”, “inframe_insertion”, “missense_variant”, “start_lost”, “stop_lost”, “protein_altering_variant”

**Synonymous**: “synonymous_variant”, “stop_retained_variant”, “incomplete_terminal_codon_variant”

To exclude common variants and to limit our analyses to biologically variant categories, the remaining calls were restricted to missense and lof variants with an allele frequency less than 0.001 in ExAC (Lek et al., 2016) that were bioinformatically predicted to be deleterious by both CADD (CADD-phred >20) (Kircher et al., 2014) and DANN (>0.99) (Quang et al., 2015). ClinVar (ftp://ftp.ncbi.nlm.nih.gov/pub/clinvar/vcf_GRCh37) and COSMIC coding mutations (https://cancer.sanger.ac.uk/cosmic/download) were used to further refine the call set to disease relevant variants.

### SNV characterization and prioritization

We used a series of gene-level and variant-level filters to identify SNVs of particular interest to human disease, as described in the main manuscript text. These filters are derived from publicly-available databases and bioinformatic prediction algorithms and represented in the columns of Table S4 and detailed below.

#### VARIANT DETAILS

**gene**, HUGO gene nomenclature committee (HGNC) official human gene symbol;

**transcript**, Ensembl transcript identifier;

**locus**, genomic coordinates from hg19 reference, and base of the reference and alternate allele;

**chr**, human chromosome on which the variant is located;

**pos**, chromosomal position of the variant in hg19 coordinates;

**ref**, reference base at this chromosomal position;

**alt**, alternate base at this chromosomal position;

**duplicated_locus**, TRUE if the site is multiallelic and has been split.

**rsid**, reference SNP identifier from dbSNP (https://www.ncbi.nlm.nih.gov/snp/);

**worst_csq**, worst consequence predicted by VEP

**consequence**, consequence predicted by VEP

**hgvsp**, Ensembl identifier of the affected protein and most likely amino acid change;

**hgvsc**, Ensembl identifier of the affected transcript and coding change;

**homs**, file name(s) of hESC line(s) in which the variant was identified in a homozygous state;

**hom_count**, number of homozygous variant among the 143 analyzed hESC lines;

**hets**, file name(s) of hESC line(s) in which the variant was identified in a heterozygous state;

**het_count**, number of heterozygous variant among the 143 analyzed hESC lines;

**AC**, total number of variant alleles genotyped among the 143 analyzed hESC lines;

**AN**, total number of alleles at that genomic location genotyped among the 143 analyzed hESC lines;

**AF**, ratio of variant to total alleles in hESCs;

#### VARIANT-LEVEL FILTERS

**gnomad_global_AC**, number of times this variant was present in 15,708 whole genome sequences currently represented in the gnomAD (https://gnomad.broadinstitute.org/) database (Karczewski et al., 2019);

**gnomad_global_AF**, ratio of variant to total alleles among the 15,708 whole genome sequences in gnomAD;

**exome_global_AC**, number of times this variant was present in the ExAC database in 60,706 exomes that are non-overlapping with the whole genomes represented in gnomAD (Lek et al., 2016);

**exome_global_AF**, ratio of variant to total alleles at this genomic location in ExAC;

**cadd_raw**, raw score from the Combined Annotation Dependent Depletion (CADD) predictor of variant deleteriousness (Kircher et al., 2014);

**cadd_phred**, scaled phred-like CADD score where the bottom 90% of deleterious variants have a score of 0-10, the next 9% have scores of 10-20, and so on;

**dann**, score of variant deleteriousness from the Deleterious Annotation of genetic variants using Neural Networks (DANN) with scores ranging from 0 to 1 for neutral to most deleterious (Quang et al., 2015);

**fathmm_xf_cod_score**, score of likely coding variant pathogenicity from the Functional Analysis Through Hidden Markov Models (FATHMM-XF) predictor (Rogers et al., 2018)

**fathmm_xf_nc_score**, FATHMM prediction for non-coding (e.g. splice donor) variants;

**fathmm_warn**, variant annotation from FATHMM at default sensitivity and specificity thresholds;

**fathmm_cancer**, FATHMM prediction of cancer-associated coding variants (http://fathmm.biocompute.org.uk/cancer.html) with annotations at default sensitivity and specificity thresholds (Shihab et al., 2013)

**cosmic_id**, Catalogue Of Somatic Mutations in Cancer (COSMIC) numerical internal database identifier (http://cancer.sanger.ac.uk/census/) (Tate et al., 2019);

**cosmic_count**, number of times the specific variant was reported in COSMIC, where multiple entries denote multiple splice isoforms;

**cosmic_cds**, base changes for each of the major splice isoforms of the protein in COSMIC,

**cosmic_aa**, amino acid changes for each of the major splice isoforms of the protein in COSMIC,

#### GENE-LEVEL FILTERS

**clin_def**, clinical syndrome(s) associated with defects in the affected gene as reported in the manually-curated ClinVar database (www.ncbi.nlm.nih.gov/clinvar/) (Landrum et al., 2014);

**clin_sig**, likely clinical significance of the variant based on manual curation of the strength of supporting data in the literature;

**clin_vc**, variant type;

**clin_db**, links to databases describing clinical syndrome(s) in greater detail;

**MIM**, identifier for the clinical syndrome(s) associated with defects the affected gene as reported in the manually-curated Online Mendelian Inheritance in Man database (http://www.omim.org/downloads) (OMIM, 2019)

**Genomic.Location**, genomic location of the gene associated with the clinical syndrome(s) described in OMIM;

**Haploinsufficiency.Description**, annotation of the gene as dosage-sensitive in the manually-curated ClinGen database (http://www.ncbi.nlm.nih.gov/projects/dbvar/clingen/) (Rehm et al., 2015)

**Loss.phenotype.OMIM.ID**, OMIM entries for genes in which gene loss of function is associated with a clinical phenotype;

**Dosage**, annotation of haploinsufficiency among the 59 genes designated by the American College of Medical Genetics (ACMG) manual (https://www.ncbi.nlm.nih.gov/projects/dbvar/clingen/acmg.shtml),

**Fertility_Related**, genes described in the literature to be related to fertility (O’Flynn O’Brien et al., 2010; Venkatesh et al., 2014);

**Imprinted**, inclusion of the gene in a manually-curated list of imprinted human genes (http://www.geneimprint.com/site/genes-by-species);

**X-linked dominant**, OMIM disease genes that show X-linked dominant inheritance according to (Berg et al., 2013)

**Tumor Suppressors**, inclusion of the gene in a curated list of tumor suppressors (https://bioinfo.uth.edu/TSGene/) (Zhao et al., 2016);

**COSMIC_tier**, inclusion of the gene as a COSMIC Tier 1 gene that has an established association to cancer (http://cancer.sanger.ac.uk/census/);

**P53_Pathway**, inclusion of the gene in the KEGG p53 pathway (https://www.genome.jp/dbget-bin/www_bget?pathway+hsa04115) (Kanehisa et al., 2017, 2019)

**Growth restricting**, Top 50 growth restricting genes in hESCs identified by (Yilmaz et al., 2018)

**CONDITION**, clinical syndrome associated with the gene in the Clinical Genomic Database (CGD, https://research.nhgri.nih.gov/CGD/);

**INHERITANCE**, reported inheritance of the clinical syndrome associated with the gene in CGD;

**COMMENTS**, comments associated with the clinical syndrome associated with the gene in CGD;

**INTERVENTION.RATIONALE**, clinical intervention rationale for the clinical syndrome associated with the gene in CGD;

**REFERENCS**, references associated with the clinical syndrome associated with the gene in CGD;

**Autosomal Dominant**, OMIM disease genes that show dominant inheritance according to (Berg et al., 2013)

**obs_mis**, observed number of missense variants in this gene in gnomAD (Karczewski et al., 2019);

**exp_mis**, expected number of missense variants in this gene in gnomAD;

**mis_z**, ExAC score of gene constraint to missense variants where positive scores indicated increased constraint (Lek et al., 2016);

**oe_mis_lower**, 90% confidence interval for the lower bound of observed to expected missense variants in this gene in gnomAD;

**oe_mis_upper**, 90% confidence interval for the upper bound of observed to expected missense variants in this gene in gnomAD;

**obs_lof**, observed number of loss-of-function variants in a gene in gnomAD;

**exp_lof**, expected number of loss-of-function variants in a gene in gnomAD;

**pLI**, probability of loss intolerance from a LoF mutation from the ExAC database based on expected versus observed LoF mutations;

**oe_lof**, mean fraction of observed to expected loss-of-function variants in a given gene;

**oe_lof_lower**, 90% confidence interval for the lower bound of observed to expected loss-of-function variants in this gene in gnomAD;

**oe_lof_upper**, 90% confidence interval for the upper bound of observed to expected loss-of-function variants (LOEUF) in this gene in gnomAD;

**constraint_flag**, Flags assigned to constrained genes as defined in gnomad v2.1 (see https://gnomad.broadinstitute.org/faq)

**gene_type**, Type of constrained gene as per gnomad v2.1 (see https://gnomad.broadinstitute.org/faq)

